# Elevated CO_2_ effects on morphology and survival rate in intertidal benthic foraminifera

**DOI:** 10.1101/843086

**Authors:** Gunasekaran Kannan, Saravanakumar Ayyappan, Karthikeyan Perumal

## Abstract

Oceans are absorbing about one-third of the anthropogenic CO_2_ from the atmosphere, and the excess CO_2_ leads to reduce the seawater pH, carbonate ion concentrations and saturation states of biologically important calcium carbonate minerals. The ocean acidification affects many calcifying organisms such as foraminifera. The present study assessed experimentally modified ocean pH impacts on survival rate and shell morphology variation in three species of intertidal benthic foraminifera. Foraminifera was collected from Parangipettai coastal waters, Tamilnadu, India. Foraminiferal specimens were cultured for a period of five weeks at three different pH treatments that replicated future scenarios of a high CO_2_ and low pH. The experimental results revealed that reduction of seawater pH significantly affected foraminiferal survival and morphology. Scanning Electron Microscopic observations revealed significant changes in foraminiferal morphology with clear evidence of dissolution and cracking processes on the test surface, and reduction in teeth structure in treatments with decreasing pH. Hence, altering the seawater chemistry might have extensive impacts on benthic foraminifera.

## 1. Introduction

The Ocean absorbs the anthropogenic CO_2_ from the atmosphere and it changes the ocean’s fundamental chemistry (Orr, 2011). The pH in ocean surface water has decreased by 0.1 units since the pre-Industrial Revolution and is now expected to drop further up to 0.4 units by the end of the 21st century (Calderia and Wicket, 2005). The atmospheric CO_2_ concentration level of 280 ppm in the beginning of industrial revaluation increased to 393 ppm in 2012 (Maona Loa Observatory, 2012) and the level is currently increasing at the rate of 0.5 % per year (Forster et al., 2007). In this industrial era, oceans have absorbed approximately 30 % of CO_2_ from the atmosphere due to the burning of fossil fuels and land use change (Sabine et al., 2004; Raven et al., 2005; IPCC, 2014). The observed CO_2_ leads to reduce the seawater pH and carbonate ion concentration, and this process is known as ocean acidification (Calderia and Wicket, 2003).

Ocean acidification is likely to have a negative effect on calcifying marine organisms such as mollusks corals, some microalgae, foraminifera, and calcifying algae. Shells, skeletons, and other protective structures of the marine organisms are mostly made up of calcium carbonate materials (CaCO_3_) (Zeebe and Sanyal, 2002; Riebesell and Tortall, 2011). Aragonite, calcite and magnesium calcite (Mg-calcite) are the three-major biogenic CaCO_3_ occurring in the seawater (Gattuso and Hansson, 2011). The concentration of carbonate ions and the CaCO_3_ saturation state are controlled the calcium carbonate precipitation where decrease processes with increasing the ocean acidification (Gattuso and Hansson, 2011).The response of marine calcifiers to decreasing calcium carbonate saturation state will be species-specific and depend on environmental parameters such as light, temperature, and available nutrients, the form of carbonate mineralogy and the mechanism of calcification (Feely et al., 2004) Although the chemistry of ocean acidification is well understood, and its impact on marine organisms and ecosystem remains in some cases poorly known (Gattuso et al., 2010). The changing seawater carbonate chemistry directly affects calcareous skeletal structures of the marine organisms (Fabry et al., 2008), and the high *p*CO_2_ treatment reduces calcification rate in benthic foraminifera (Kuroyanagi et al., 2009; Dissard et al., 2010; Fujita et al., 2011; Haynert et al., 2011; Uthicke et al., 2013).

Foraminifera is one of the most abundant groups of calcifying organisms, approximately precipitate Ca. 50% of biogenic calcium carbonate in the open oceans (Faber and Preisig, 1994; Schiebel, 2002). Most of the foraminifera test walls are organic, agglutinated or made of calcium carbonate in various chemical and structural modifications (Hemleben et al., 1986). Calcium carbonate forms are composed of either magnesium calcite, calcite or aragonite (Hemleben et al., 1986). Approximately 900 described genera deposit tests of calcium carbonate (Lipps, 1973). Foraminifera shells are composed of primary and secondary layers of calcite (Erez, 2003), primary layer consists of high-Mg calcite (Bentov and Erez, 2006) and the second layer consists of low-Mg calcite (Haynert et al., 2011). Calcite producing foraminifera are one of the most important groups of calcite producers. They precipitate annually 1.4 billion tons of calcite, which accounts for 25% of the total global calcite production (Langer, 2008).

Foraminifera are an interesting group to examine in relation to ocean acidification experiments, they distributed worldwide, are environmentally sensitive, have short life-history and they are important ecosystem engineers (Jones et al., 1994). Some foraminifera species showed sensitivity to ecological conditions (Faber and Preisig, 1994). The most of the ocean acidification research on foraminifera has been published in the last 8 years with a variety of responses such as increased shell weight with increased carbonate ion concentration (Bijma et al., 1999, 2002), diversity reduction and species abundance shift from 24 to 4 species in natural pH levels (Dias et al., 2010), high pCO_2_ site impact many foraminifera tests with corroded or pitted (Fabricius et al., 2011).

In recent years, the atmospheric *p*CO_2_ levels have increased in the southwest Bay of Bengal (Shanthi et al., 2016). CO_2_ flux indicates that the southeast Bay of Bengal acts as a significant sink of CO_2_ from the atmosphere (Singh and Ramesh, 2015). In the present study area of coastal Parangipettai in southeast Bay of Bengal, the *p*CO_2_ level ranged from 377 to 390 µatm and seawater *p*CO_2_ levels ranged from 156 to 980 µatm. The present study investigated the survival and morphology of benthic foraminifera *Ammonia beccarii, A. tepida* and *A. dentata* in three different pH treatments that replicated future scenarios of a high atmospheric CO_2_ and low pH.

## 2. Materials and methods

### 2.1 Sample Collection

The surface sediments scrapes (1-2 cm depth) containing benthic foraminifera were collected from the Parangipettai coastal waters, Bay of Bengal, Tamilnadu, India (11º 30’ 45. 06’’ N and 79º 46’ 29.18’’ E) during low tide in April 2016. The sediment sample was transferred to the laboratory, and then they were mixed and sieved over a set of 63 µm and 1000 µm screens.

### 2.2 Foraminiferal isolation

By simple observations of a small amount of sediment through a compound binocular microscope, live specimens of *A. beccarii, A. dentata* and *A. tepida* were identified. Those individuals showed colorful protoplasm extensively distributed across the entire foraminiferal tests, except in the last chamber. Despite the naturally intense protoplasm color is widely used as an indicator of the viable foraminiferal individuals (Bernhard 2000), pseudopodia activity was observed to confirm the viability. The procedure of sampling sieving and observation of live specimens was repeated for two days until a large number of target foraminifera was achieved (10 −15 specimens/ cm^3^ of wet sediment). Approximately 100 cm^3^ of mixed sediment was placed in a series of 500 cm^3^ filtering flasks, and these glass containers were filled up with filtered natural seawater with ~ 33 ppt of salinity. Finally, 4500 living specimens were randomly selected and transferred to foraminiferal culture chambers. Experiment duration was a five-week period, and during the acclimation time the pH was reduced gradually until each treatment reached its required pH levels/*p*CO_2_. Probes measuring pH and temperature continually monitored four reservoir tanks.

### 2.3 Experimental setup

The seawater pH was continually manipulated by bubbling air with a known equivalent atmospheric concentration of CO_2_ (approx. 380, 1000 and >1200 µatm) into three reservoir tanks, separately. Therefore, the three selected pH levels of 8.1, 7.7 and 7.6 representing the range of pH predicted for future scenarios in a high CO_2_ level. Seawater was continuously pumped from the 400 L tanks into culturing chambers.

Seawater samples were taken from each mesocosm tank to measure the total alkalinity (A_T_). Collected seawater samples were stored in borosilicate glass bottles (250 ml) and it transferred to the laboratory. The total alkalinity concentration in seawater was analyzed by using an automatic alkalinity titration kit (Metrohm, Switzerland) at room temperature (29ºC). The measured values of salinity, temperature, pH and total alkalinity (A_T_) were used to calculate the carbon system parameters such as dissolved inorganic carbon (DIC) *p*CO_2_, bicarbonate ions (HCO_3_^−^) carbonate concentration (CO_3_^2−^) and saturation states of calcite (Ω_Calcite_) and aragonite (Ω_Aragonite_) version of the program CO_2_ Cal.

After completing the experimental period, all chambers were opened up and inserts were removed and placed to a petri dish. All the foraminiferal samples were fixed and stained immediately. Foraminifera was alive at the end of the experiment. Rose Bengal is a biological stain that absorbs onto proteins which are major cytoplasmic components and stains them rose color (Bernhard 2000). Ten live specimens with intact tests from each treatment were selected for SEM analysis after the morphological observation of stereomicroscopy. The specimens were mounted onto SEM tubs using double-sided adhesive tabs and subsequently, were imaged with a JSM 5610LV.

## 3. Results

### 3.1 Seawater carbonate chemistry

Seawater temperature (29 ºC) and salinity (33 ppt) were stable during the incubation period. As a result of three different *p*CO_2_ treatments, the pH values ranged from 8.10 to 7.6 (Table 2). The seawater total alkalinity ranged from 2240 to 2290 A_T_ µmol kg^−1^. Bicarbonate levels increased with increasing pCO2 levels. The higher *p*CO_2_ treatment reduced the availability of carbonate ion in experimental seawater. The important calcite Ω_calc_ and aragonite Ω_ar_ minerals reduced with increasing levels of the *p*CO_2_ (Table 2).

### 3.2 Survival rate

From a total of 4502 specimens transferred into the culturing chambers, only 3973 specimens were retrieved at the end of the experiment in all three species. This indicated a loss of 529 individuals throughout the experimental period (Table 1 and Fig. 1). This is attributed to the loss of specimens while transferring as well as the migration of specimens out the culturing chambers. Survival rate was calculated as a percentage of the total number of surviving individuals compared with the total number of retrieved individuals at the end of the experiment.

**Figure 1.**
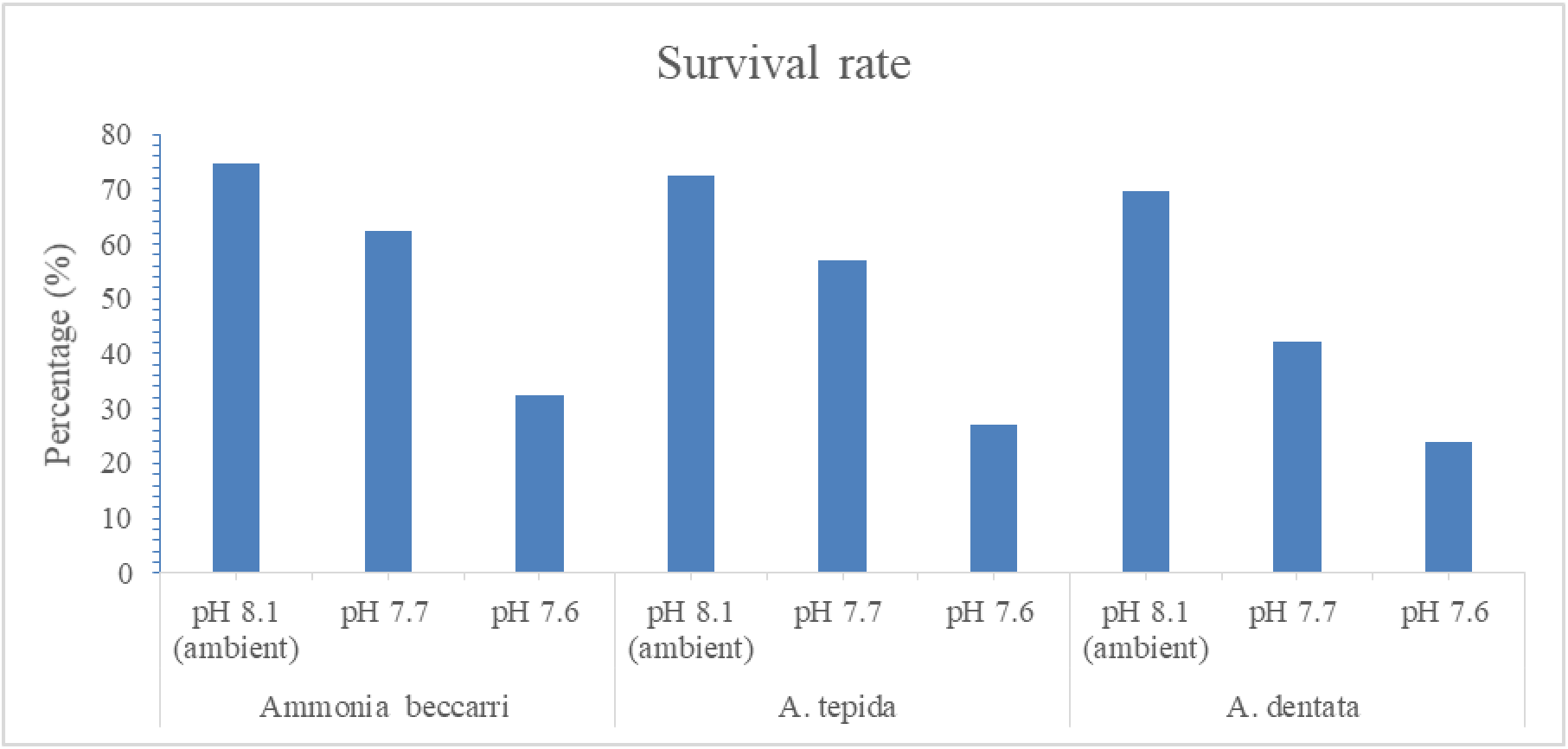
Survival rate of foraminifera specimens (*Ammonia beccarii, A. tepida and A. dentata*) in ambient and low pH treatment.

**Table 1.**
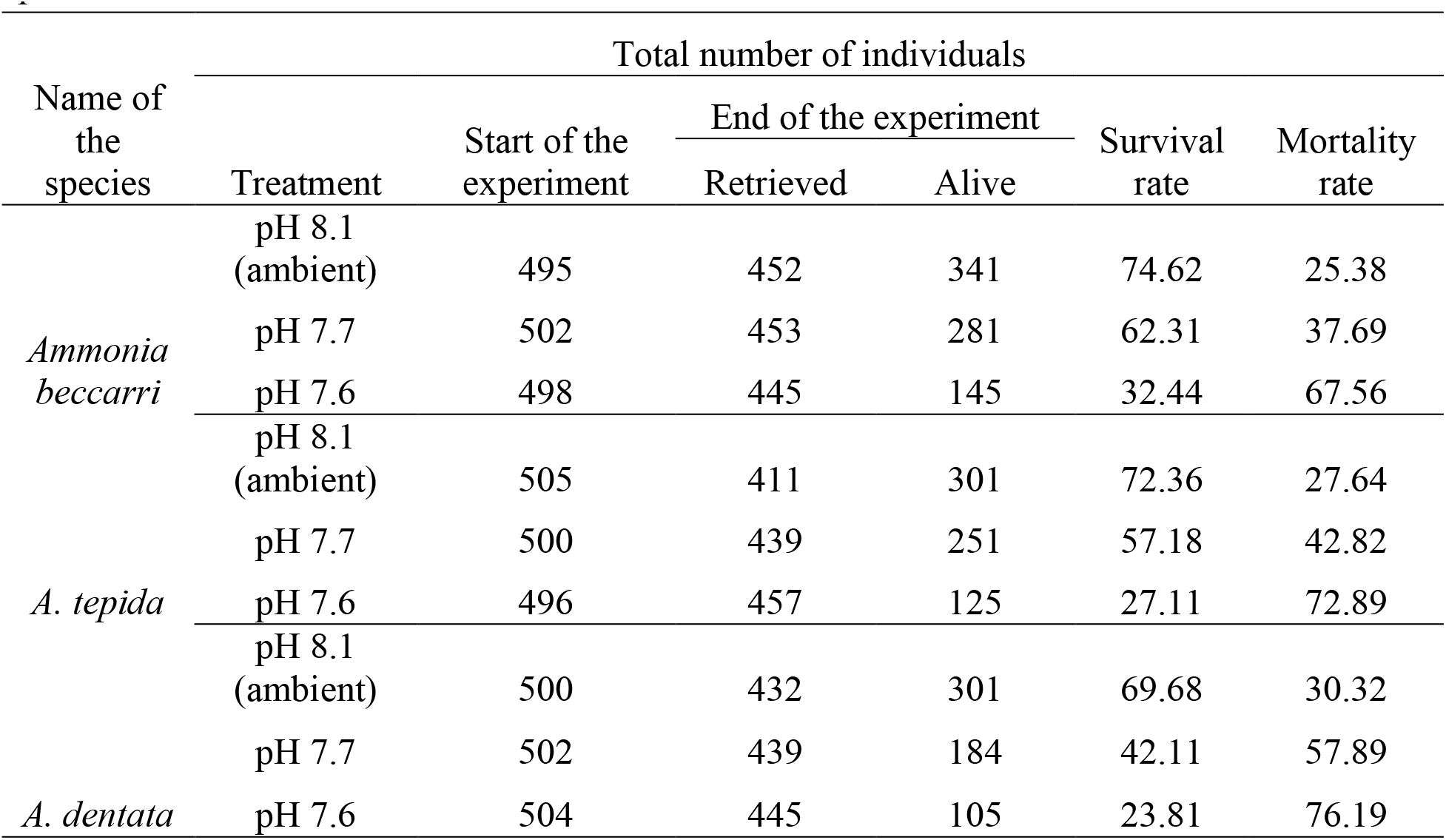
Number of individuals of *Ammonia beccarri A. tepida* and *A. tendata* cultured under different pH conditions ambient pH 8.1, pH 7.7 and pH 7.6. Individuals showing one or more new chambers added during the experimental period were considered as live individuals, survival rate (%) was calculated based on the number of living and dead over the experimental period.

**Table 2.** The low pH experiment impacts on foraminifera individuals effects

| Experiment finding on test morphology foraminifera species |  |  |  |
| --- | --- | --- | --- |
| Treatment | *Ammonia beccarii* | *A. tepida* | *A. dentate* |
| pH 8.1 (Ambient) | no affects | no affects found | Effects Not found |
| pH 7.7 | No affects | ventral side of notch slightly dissolved | Folium part of the umbilical side cracked |
| pH 7.6 | surface ornamentation dissolute | surface cracking found | dorsal and ventral side test dissolved |

Foraminifera culture chamber was opened and specimens were categorized as live or dead based on the rose Bengal staining. Any specimens that were inactive or contained no cytoplasm were considered as dead. The low survival rate was observed at pH 7.6 (Fig.1), and it was least in *A. dentata* among the three species. The highest survival rate was observed at ambient pH and pH 7.7 in all three species. The moderate survival rate was observed in pH of 7.6.

### 3.3 Scanning Electron micrographs (SEM) of foraminiferal tests

SEM images of all three *A. beccarii, A. tepida*, and *A. dendata* showed morphological differences among specimens cultured at different pH conditions (Fig. 2). These observations indicated a progressive alteration of the foraminiferal morphology. The test surface was smooth, and the pore size and chamber arrangements remained unaffected in all the three species of foraminifera at ambient pH 8.10. The most significant features observed on the test surface appearance were cracking and dissolution on the individuals exposed to the lowest pH levels. The foraminiferal specimens that were cultured at pH 7.7 and 7.6 showed the same cracking and dissolution of the surface ornamentation. The reduced pH treatments affected the foraminiferal folium part of the umbilical side and dissolved the chamber suture. More importantly, *A. dentata* was observed to be moderately affected in two pH treatments on whorl suture and dissolved test surface, and the cracked chamber was also found on the dorsal side of the species (Fig.2).

**Figure 2.**
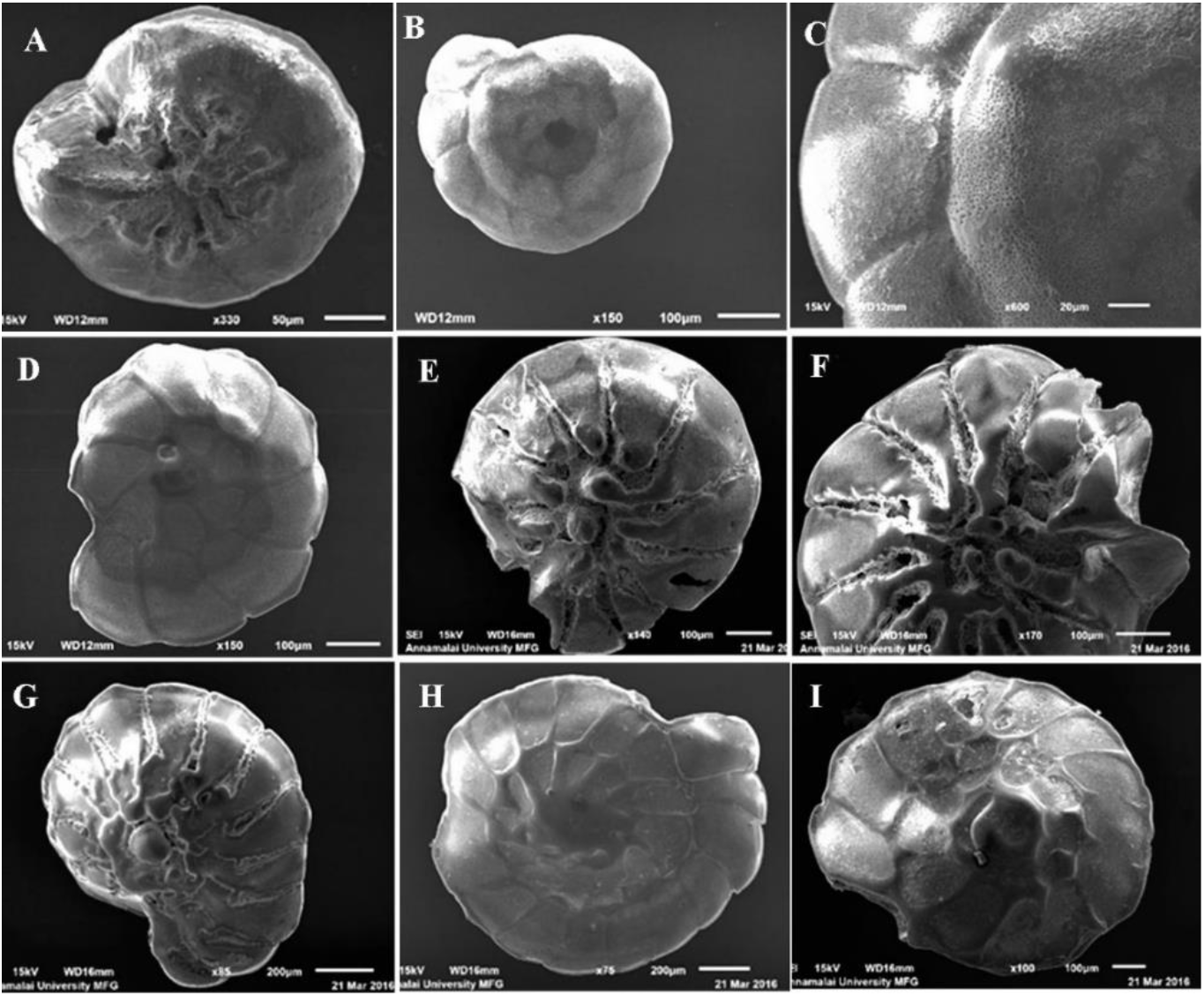
SEM micrographs of benthic foraminifera in three different pH ambient pH 8.10 pH 7.7 and pH 7.6 experiment. *A. beccarii* in Ambient pH 8.10 pH 7.7 and pH 7.6 no variation found (A, B, D, and G), surface dissolute (C), *A.tepida* pH 7.6, Ventral side dissolute and test cracked (E and F), pH 7.7 and pH 7.6 dorsal and ventral side test dissolved (H and I) pH 7.7 and pH 7.6.

## 4. Discussion

The seawater carbonate ions and calcite are important carbon species to make the test on skeleton of calcifying organisms (Keul et al., 2013). The experimental seawater bicarbonate concentration increases with increasing *p*CO_2_, in contrast, the carbonate iron and mineral calcite reduces with increasing *p*CO_2_.

### 4.1 Survival rate

The present experiment was conducted to determine the short-term responses of three benthic foraminiferal species (*Ammonia beccarii, A. tepida* and *A. dentata*) to three pH treatments. All the pH treatments caused high mortality in the species due to negative effect of reduced pH and also to the sudden exposure to experimental conditions from natural environment. The mortality was well pronounced in *A. dentata* at pH 7.6, as compared to other two species, and this reveals species-specific vulnerability to change in pH, in support of earlier works (Fabry et al., 2008; Keul et al., 2013). The lower pH did not significantly affect the survival rate in agglutinated and tectinous benthic foraminifera but negatively impacted the survival rate in calcareous foraminifera (Bernhard et al., 2009a). The present study recorded noticeable mortality rates in all the three species with decreasing pH, similarly to the previous work of, Mclntyre Wressing et al. (2011) who have reported that higher CO_2_ concentration affects survival rate in *Amphistegina gibbosa*.

### 4.2 Scanning electron micrograph of foraminifera

SEM analysis demonstrated that the morphology of all three species were sensitive to decreasing pH conditions. In the ambient pH condition, the morphology of all three species were very smooth with intact ornamentation such as tubercles and teeth lining the aperture. The present study observed morphological variation such as, dissolution in ventral side of notch and cracking in umbilical part of *A. tepida* and *A. dentata* pH 7.7 treatment, but there was no such changes found in *A. beccarii* at 7.7 treatment. Khana et al. (2013) have recorded that shell appearance of *Haynesina germanica* is not damaged at 586 ppm of CO_2_ treatment as well as teeth shape and size are intact, but only teeth numbers are reduced. However, they have found that the average number of teeth, length and width of *H. germanica* teeth structure are reduced at 750 ppm of CO_2_ treatment (Khana et al., 2013). Haynert et al. (2011) have reported that the test surface of *Ammonia aomoriensis* is not affected when treated between 618 to 751 µatm. The present study recorded broken teeth with dissolved test surface to confirm the adverse effect of lower pH on morphology of foraminiferans. A similar observation has been made in gastropods with reduced toughness of inner shell at low pH of 7.7 (Duquette et al., 2017) and also with reduced shell weight due to shell dissolution with reduced pH and increased CO_2_ (Nienhuis et al., 2016). The present study also recorded higher test weight loss in *A. dentata* in pH 7.7 and pH 7.6 treatments.

The results revealed that all the three foraminiferal species when exposed to the higher CO_2_ (pH 7.6) exhibited morphological deformations such as dissolution, rounding teeth, surface cracking, dorsal and ventral side test dissolved, and these deformations might have affected the survival of ferminiferans. This is accordance with White et al. (2013) who have reported that high CO_2_ exposure results in smaller shells and reduces the larval survival rate in *Argopecten irradiants*

Similar to our observations, Haynert et al. (2011) have found that 1 to 3 younger chambers of *A. aomoriensis* are severely decalcified at *p*CO_2_ levels 929 to 1829 µatm, in addition to cracks found around the pores in the treatment. The authors also have recorded that increasing high *p*CO_2_ of 3130 µatm treatment reduces test diameter, the growth rate of *A. aomoriensis* and destroys all chambers by complete calcium carbonate dissolution (Haynert et al., 2011). Riebsell et al. (2000) have found that the increasing CO_2_ concentration affects the coccoliths in term of malformed coccoliths, increases incomplete coccospheres, and reduces calcification rate by 36 to 83 %. This work has revealed that cultured coccolithophorids are highly sensitive to increased seawater *p*CO_2_. Keul et al. (2013) have reported that *growth* rate in *Ammonia* species decreases with decreasing [CO^2-^_3_] Ω_cacl_< 1. The present observations are in support of previous laboratory studies, where *A. aomoriensis* and *A. beccarii* are affected by test degradation (Le Cade, 2003; Haynert et al., 2011). Therefore, CO_2_ induced ocean acidification experimental studies reiterated the vulnerability of the benthic foraminifera to near future ocean acidification, by causing morphological deformation and absence of important feeding structures in the organisms.

## 5. Conclusion

The experimental study provides a detailed understanding of the negative effects on survival rate, shell morphological changes in the benthic foraminifera in low pH experiment. The changes in the carbonate chemistry of natural coastal environment will have similar effects to this simulated in laboratory condition and hence, calcifying organisms will be more vulnerable to the near future ocean acidification, as reiterated by the present study.

## 6. Acknowledgments

The authors are thankful to Dean and Director for given the valuable support during the study period and to the DST – INSPIRE programme for financial assistance to Gunasekaran (IF130109).

## References

Bentov, S., and Eewz, J., (2006). Impact of biomineralization processes on the Mg content of foraminiferal shells: A biological perspective. Geochem. Geophys. Geosyst, 7, Q01P08.

Bernhard, J. M., (2000). Distinguishing live from dead foraminifera: Methods review and proper applications. Micropaleontol.46, 38–46.

Bernhard, J. M., Barry, J., Buck, K., and Stanczak, V. (2009a). Impact of intentionally injected carbon dioxide hydrate on deep sea benthic foraminiferal survival. Global Change Biol., 15, 2078–2088.

Bijma, J., Honisch, B. and Zeebe, R.E. (2002). Impact of the ocean carbonate chemistry on living foraminiferal shell weight: comment on “Carbonate ion concentration in glacial-age deep waters of the Caribbean Sea” by W.S. Broecker and E. Clark, Geochem. Geophy. Geosy. 3, 1064.

Bijma, J., Spero, H.J. and Lea, D.W. (1999). Reassessing foraminiferal stable isotope geochemistry: Impact of the oceanic carbonate system (experimental results), in: Fischer, G., Wefer, G. (Eds), Use of Proxies in Paleoceanography: Examples from the South Atlantic. Springer-Verlag. New York. 489–512.

Caldeira, K. and Wickett, M.E. (2003). Anthropogenic carbon and ocean pH. Nature, 425, 325.

Caldeira, K. and Wickett, M.E. (2005). Ocean model predictions of chemistry changes from carbon dioxide emissions to the atmosphere and ocean. J. Geophys. Res.110.

Dias, B.B., Hart, M.B., Smart, C.W., and Hall-Spencer, J.M. (2010). Modern seawater acidification: The response of foraminifera to high-CO2 conditions in the Mediterranean Sea. J. Geol. Soc.167, 843–846.

Dissard, D., Nehrke, G., Reichart, G.J. and Bijma, J. (2010). Impact of seawater pCO2 on calcification and Mg/Ca and Sr/Ca ratios in benthic foraminifera calcite: results from culturing experiments with *Ammonia tepida*. Biogeosciences, 7, 81–93.

Duquette, A., McClintock, J.B., Amsler, C.D., Pérez-Huerta, A., Milazzo, M. and Hall-Spencer, J.M. (2017). Effects of ocean acidification on the shells of four Mediterranean gastropod species near a CO_2_ seep.124, 917–928.

Erez, J. (2003). The Source of Ions for Biomineralization in Foraminifera and Their Implications for Paleoceanographic Proxies. Rev. Mineral. Geochem, 54, 115–149.

Faber, W.W. and Preisig, H.R. (1994). Calcified structures and calcification in protists. Protoplasma, 181, 78–105.

Fabricius, K.E., Langdon, C., Uthicke, S., Humphrey, C., Noonan, S., De’ath, G., Kazaki, R., Muehllehner, N., Glas, M.S. and Lough, J.M. (2011). Losers and winners in coral reefs acclimatized to elevated carbon dioxide concentrations. Nat. Clim. Change, 1, 165–169.

Fabry, V.J., Siebel, B.A., Feely, R.A. and Orr, J.C. (2008). Impacts of ocean acidification on marine fauna and ecosystem processes. ICES J. Mar. Sci. 65, 414–432.

Feely, R.A., Sabine, C.L., Lee, K., Berelson, W., Kleypas, J., Fabry V.J. and Millero, F.J. (2004). Impact of anthropogenic CO_2_ on the CaCO_3_ system in the oceans. Science, 305, 362–366.

Forster, P., Ramaswamy, V., Artaxo, P., Berntsen, T., Betts, R., Fahey, D. W., Haywood, J., Lean, J., Lowe, D. C., Myhre, G., Nganga, J., Prinn, R., Raga, G., Schultz, M. and Dorland, R. V. (2007). Changes in atmospheric constituents and in radiative forcing. In: Solomon, S., Qin, D., Manning, M., Chen, Z., Marquis, M., Averyt, K. B., Tignor, M. & Miller, H. L. (Eds) Climate Change 2007: The Physical Science Basis. The contribution of Working Group I to the Fourth Assessment Report of the Intergovernmental Panel on Climate Change. Cambridge University Press, Cambridge.

Fujita, K., Hikami, M., Suzuki, A., Kuroyanagi, A. and Kawahata, H. (2011). Effects of ocean acidification on calcification of symbiont-bearing reef foraminifers. Biogeosciences, 8, 1809–1828.

Gattuso, J.P. and Hansson, L. (2011). Ocean acidification: background and history. In: Gattuso, J.–P. & Hansson, L. (Eds) Ocean Acidification. Oxford University Press, Oxford. pp. 1–20.

Gattuso, J.P., Gao, K., Lee. K., Rost, B. and Schultz, K.G. (2010). Approaches and tools to manipulate the carbonate chemistry. In: Riebesell, U., Fabry, V. J., Hansson, L. & Gattuso, and J.-P. (Eds) Guide to Best Practices for Ocean Acidification Research and Data Reporting. Publications Office of the European Union, Luxembourg, pp. 41–52.

Haynert, K., Schönfeld, J., Riebesell, U. and Polovodova, I. (2011). Biometry and dissolution features of the benthic foraminifer *Ammonia aomoriensis* at high pCO_2_. Mar. Ecol. Prog. Ser.432, 53–67.

Haynes, J.R. (1981). Foraminifera. Macmillan Publishers Ltd, London.

Hemleben, C.H., Anderson, R.O., Berthold, W. and Spindler, M. (1986). Calcification and chamber formation in Foraminifera-a brief overview. In: Leadbeater, B. S. C. and Riding, R. (Eds) Biomineralization in Lower Plants and Animals. The Systematics Association Special Volume No. 30. Oxford University Press, Oxford. Pp.237–249.

IPCC, 2014: Summary for Policymakers. In: Climate Change 2014: Mitigation of Climate Change. The contribution of Working Group III to the Fifth Assessment Report of the Intergovernmental Panel on Climate Change [Edenhofer, O., R. Pichs-Madruga, Y. Sokona, E. Farahani, S. Kadner, K. Seyboth, A. Adler, I. Baum, S. Brunner, P. Eickemeier, B. Kriemann, J. Savolainen, S. Schlömer, C. von Stechow, T. Zwickel and J.C. Minx (eds.)]. Cambridge University Press, Cambridge, United Kingdom and New York, NY, USA.

Jones, C.G., Lawton, J. H. and Shachak, M. (1994). Organisms as ecosystem engineers. Oikos, 69, 373–386.

Keul, N., Langer, G., de Noojier, L. J. and Bijma, J. (2013). Effect of ocean acidification on the benthic foraminifera *Ammonia* sp. is caused by a decrease in carbonate ion concentration. Biogeosciences, 10, 6185–6198.

Khana, K., Godbold, J.A., Austin, W.N. and Peterson, D.M. (2013). Impact of ocean acidification on the functional morphology of foraminifera. Plos one 8, 1–4.

Kuroyanagi, A., Kawahata, H., Suzuki, A., Fujita, K. and Irie, T. (2009). Impacts of ocean acidification on large benthic foraminifers: results from laboratory experiments. Mar. Micropaleontol.73, 190–195.

Langer, M.R. (2008). Assessing the contribution of foraminifreran protists to global ocean carbonate production. J. Eukaryot. Microbiol. 55, 163–169.

Le Cadre, V., Debenay J. P. and Lesourd, M. (2003). Low pH effects on *Ammonia beccarii* test deformation: implications for using test deformations as a pollution indicator. J. Foraminiferal Res. 33, 1 - 9.

Lipps, J.H. (1973). Test Structure in Foraminifera. Annu. Rev. Microbiol., 27, 471–486.

Maona Loa Observatory, (2012). Hawaii’s Mauna Loa Observatory: Fifty Years of Monitoring the Atmosphere.

Mclntyre-Wressnig, A., Bernhard, J. M., McCorkle, D.C. and Hallock, P. (2011). Non-lethal effects of ocean acidification on two symbiont-bearing benthic foraminiferal species. Biogeosciences Discuss., 8, 9165–9200.

Nienhuis, S., Palmer, A.R. and Harley, C.D.G. (2016). Elevated CO_2_ affects shell dissolution rate but not calcification rate in a marine snail. Proc. R. Soc. B., 277, 2553–2558.

Orr, J.C. (2011). Recent and future changes in ocean carbonate chemistry. In: Gattuso, J.P and L. Hansson, (Eds) Ocean acidification. Oxford University Press, Oxford. Pp. 41–66.

Raven, J., Caldeira, K., Ove Hoegh, G., Liss, P., Riebesell, U., Shepherd, J., Turley, C. and Watson, A. (2005). Acidification due to increasing carbon dioxide. In: Policy Report 12/05. The Royal Society, London, UK, 233 pp.

Riebesell, U. and Tortell, D. (2011). Effects of ocean acidification on pelagic organisms and ecosystems. In: Gattuso, J.-P. & Hansson, L. (Eds) Ocean Acidification. Oxford University Press, Oxford. pp. 99–121.

Riebesell, U., Zondervan, I., Rost, B., Tortell, P.D., Zeebe, R.E. and Morel, F.M.M. (2000). Reduced calcification of marine plankton in response to increased atmospheric CO_2_. Nature 407, 364–367.

Sabine, C.L., Feely, R.A., Gruber, N., Key, R.M., Lee, K., Bullister, J.L., Wanninkhof, R., Wong, C.S., Wallace, D.W.R., Tilbrook, B., Millero, F.J., Peng, T.H., Kozyr, A., Ono, T. and Rios, A.F. (2004). The oceanic sink for anthropogenic CO_2_. Science, 305, 367–371.

Schiebel, R. (2002). Planktonic foraminiferal sedimentation and the marine calcite budget. Global Biogeochem. CY, 16, 1065.

Shanthi, R., Poornima, D., Naveen, M., Thangaradjou, T., Choudhry, S.B., Rao, K.H. and Dadhwal V.K. (2016). Air-sea CO_2_ flux pattern along the southern Bay of Bengal waters. Dynamics and Atmospheres and Oceans 76, 14–29.

Singh, A. and Ramesh, R. (2015). Environment controls on new and primary production in the northern Indian Ocean. Progress in Oceanography 131, 138–145.

Uthicke, S., Momigliano, P. and Fabricius, K. (2013). High risk of extinction of benthic foraminifera in this century due to ocean acidification. Scientific reports. 3, 1769,

White, M.M., McCorkle, D.C., Mullineaux, L.S. and Cohen, A.L. (2013). Early exposure of bay scallop (*Argopecten irradians*) to high CO_2_ causes a decrease in larval shell growth. Plos one 8, e61065.

Zeebe, R. E. and Sanyal, A. (2002). Comparison of two potential strategies of planktonic foraminifera for house building: Mg2+ or H-removal? Geochimica ET Cosmochimica Acta, 66, 1159–1169.

